# Absence of the Spindle Assembly Checkpoint restores mitotic fidelity upon loss of sister chromatid cohesion

**DOI:** 10.1101/262204

**Authors:** Rui D. Silva, Mihailo Mirkovic, Leonardo G. Guilgur, Om S. Rathore, Rui Gonçalo Martinho, Raquel A. Oliveira

## Abstract

Sister chromatid cohesion is essential for faithful mitosis, as premature cohesion loss leads to random chromosome segregation and aneuploidy, resulting in abnormal development. To identify specific conditions capable of restoring defects associated with cohesion loss, we screened for genes whose depletion modulates *Drosophila* wing development when sister chromatid cohesion is impaired. Cohesion deficiency was induced by knock-down of the acetyltransferase Separation anxiety (San)/Naa50, a cohesin complex stabilizer. Several genes whose function impacts wing development upon cohesion loss were identified. Surprisingly, knockdown of key Spindle Assembly Checkpoint (SAC) proteins, Mad2 and Mps1, suppressed developmental defects associated with San depletion. SAC impairment upon cohesin removal, triggered by San depletion or artificial removal of the cohesin complex, prevented extensive genome shuffling, reduced segregation defects and restored cell survival. This counterintuitive phenotypic suppression was caused by an intrinsic bias for efficient chromosome bi-orientation at mitotic entry, coupled with slow engagement of error-correction reactions. We conclude that mitotic timing determines the severity of defects associated with cohesion deficiency. Therefore, although divisions are still error-prone, SAC inactivation enhances cell survival and tissue homeostasis upon cohesion loss.

## Introduction

Mitosis is a dynamic culmination of the cell cycle, consisting of rapid packaging, alignment and partitioning of the replicated chromosomes. This process has to be tightly regulated and synchronous, as defective segregation leads to aneuploidy, a key hallmark of cancer and other human pathological conditions. The fidelity of mitosis depends on cohesive forces that keep sister chromatids together. Sister chromatid cohesion is mediated by cohesin, a tripartite ring complex that embraces sister chromatid fibers from the time of their replication until the subsequent mitosis (Guacci et al., 1997; Haering et al., 2008; Michaelis et al., 1997). Cleavage of cohesin by separase, a cysteine protease, marks the anaphase onset, where single chromatids are dragged to the poles by the mitotic spindle after cohesive forces are destroyed (Uhlmann et al., 1999; Uhlmann et al., 2000). Cohesin’s association with chromatin is regulated by numerous factors that ensure its loading, stability and dual mode of chromatin release (reviewed in (Mirkovic and Oliveira, 2017)). Among those, the N-terminal acetyltransferase Separation anxiety (San) (also known as Naa50) is required for establishment and/or maintenance of sister chromatid cohesion, and was recently proposed to acetylate the N-terminus of Rad21 cohesin subunit and regulate the interaction between Rad21 and Smc3 (Hou et al., 2007; Ribeiro et al., 2016; Rong et al., 2016; Williams et al., 2003). Accordingly, defects associated with San knock-down can be efficiently suppressed by several conditions that enhance cohesin stability on chromatin (Ribeiro et al., 2016).

Loss of sister chromatid cohesion is catastrophic for the cell as premature release of cohesive forces leads to random chromosome segregation. Genome randomization is highly enhanced upon cohesin loss as the presence of isolated sisters chromatids triggers continuous engagement of the error–correction machinery, resulting in extensive shuffling of chromatids between cell poles (Mirkovic et al., 2015; Oliveira et al., 2010). Error correction mechanisms are mediated by the Aurora-B kinase, which is able to sense the degree of tension across and/or between kinetochores and release erroneous chromosomal attachments that are not under tension (Foley and Kapoor, 2013; Khodjakov and Pines, 2010; Nezi and Musacchio, 2009). Upon cohesin loss, isolated chromatids lack the opposing forces to ensure proper tension and consequently undergo cycles of chromosome attachment and de-attachment, in a futile attempt to achieve chromosome biorientation. Consequently, mitosis in absence of cohesion results in random chromosome segregation, with close to absolute probability of generating aneuploid cells.

Premature release of cohesive forces during mitosis is prevented by a safeguard mechanism known as the Spindle Assembly Checkpoint (SAC) (reviewed in (Foley and Kapoor, 2013; Musacchio and Salmon, 2007). In the presence of unattached kinetochores, this safeguard mechanism prolongs mitosis by inhibiting the Anaphase Promoting Complex/Cyclosome (APC/C). SAC ensures that cohesin cleavage does not occur until all chromosomes are bioriented by blocking the APC/C, whose activation is needed for separase activity. In contrast to its known role as a safeguard mechanism for mitotic fidelity, we describe the unexpected observation that removal of the SAC alleviates mitotic errors when sister chromatid cohesion is compromised.

## Results

### *Drosophila* wing modifier screen reveals that depletion of Mad2 and Mps1 suppresses the developmental defects associated with loss of cohesion

Although the consequences of cohesin loss in unicellular organisms and cultured cells are well established, its impact on tissue proliferation and morphogenesis is poorly understood. To probe for conditions that would enhance or suppress cellular and tissue responses to cohesion defects, we performed a modifier screen in the adult *Drosophila* wing. We focused our analysis on the regulatory N-terminal acetyltransferase San, as our previous work demonstrated that knock-down of this protein during development gives rise to an intermediate adult wing phenotype that is sensitive to phenotypic modulation (Figure 1A,B) (Ribeiro et al., 2016). To search for modifiers (enhancers and suppressors) of the adult wing phenotype induced by San depletion, we co-expressed the *san* RNAi with 2955 RNAis, which theoretically depletes 2920 gene products, specifically in larvae imaginal wing discs (using the Nubbin-Gal4 driver (Brand and Perrimon, 1993; Ng et al., 1996; Wu and Cohen, 2002) (Figure 1A). The resulting wings were scored in 5 categories, according to the severity of the phenotype (Figure 1C) (Ribeiro et al., 2016). Co-expression of *san* RNAi and a control RNAi transgene did not modify the adult wing phenotype when compared to *san* RNAi transgene alone (Figure 1E) (Ribeiro et al., 2016). Any isolated enhancer gene whose depletion alone resulted in adult wings phenotypes was discarded (Fig. S1). All tested RNAi lines and scored wing phenotypes are shown in Tables S1 to S4.

**Figure 1.**
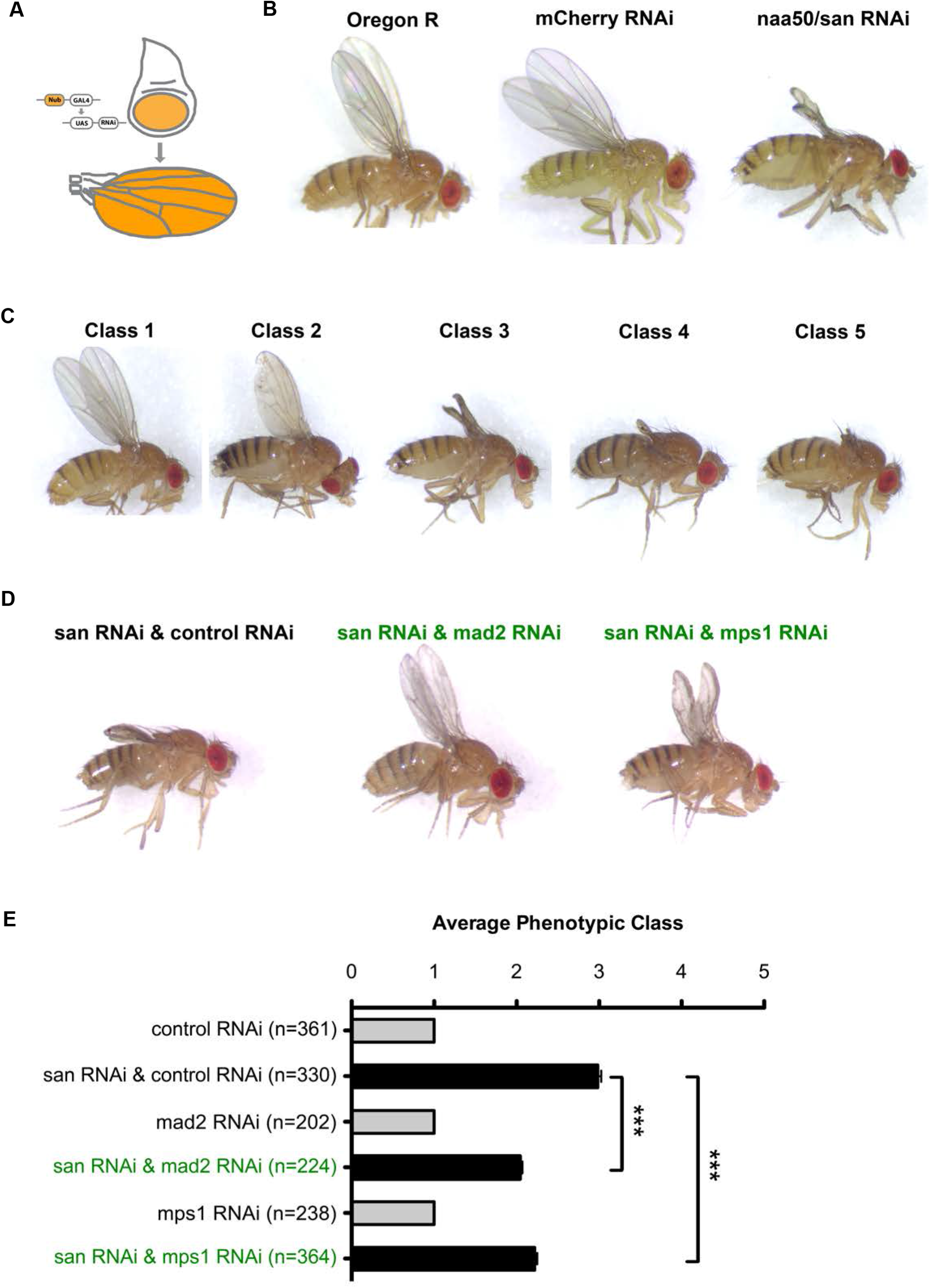
Inhibition of SAC suppresses *san* RNAi-induced adult wing developmental defects. **A.** Tissue-specific RNAi in the pouch of the larvae wing imaginal using the nubbin-Gal4 driver and the UAS/Gal4 system. **B.** Adult wings of wild type *Drosophila* (Oregon R), *Drosophila* expressing a control RNAi (*mCherry* RNAi) or expressing RNAi for san in the larvae wing imaginal discs. **C.** Adult wing phenotypic classes scored during the screen: class 1 (wild type wings); class 2 (weak wing developmental defects); class 3 (*san* RNAi-like wing phenotype); class 4 (highly abnormal wings); class 5 (absence or vestigial adult wings). Additional examples of the scored phenotypic classes are shown in (Ribeiro et al., 2016). **D.** Representative adult *Drosophila* wing phenotypes co-expressing *san* RNAi with *mCherry* RNAi, *mad2* RNAi or *mps1* RNAi in the larvae wing imaginal discs. **E)** Quantification of *Drosophila* wing phenotypes expressing individual RNAi transgenes for *mCherry, mad2* or *mps1* (grey bars) or co-expressing *san* RNAi with *mCherry* RNAi, *mad2* RNAi or *mps1* RNAi (black bars) in the larvae wing imaginal discs. Phenotypic quantification of adult wings is mean ± SD of three independent experiments and is based on the classes described in (C) (***p < 0.0001, One-way ANOVA with Bonferroni’s multiple comparison test; *n* represents the total number of flies evaluated).

We identified 19 suppressors and 10 enhancers whose depletion specifically modified the *san* RNAi adult wing phenotype (Table 1) (Figure S1). As expected, the screen revealed components previously implicated in cohesin dynamics (*Mau2* and *eco*), validating its accuracy at isolating modifiers of cohesion state (Table 1) (Ribeiro et al., 2016). This approach also identified the cohesion component *vtd*/RAD21 RNAi as a modifier of *san* RNAi adult wing phenotype (Ribeiro et al., 2016). Most of the 29 genes identified in the screen, were already characterized in *Drosophila* and/or in other species (Table 2). About half of the identified genes were either related with mitosis (*Claspin, asp, Mps1, Eb1, eco, Mau2, γTub23C* and *mad2*) or with gene expression (CG5589, *JMJD7, Pabp2, His3* and *jumu*). Other identified genes were described to be important for maintaining apicobasal cell polarity and for actin cytoskeleton organization (*capu, cno* and *Cad99C*). We identified additional suppressors/enhancer genes related with different metabolic processes (*Sfxn1-3*, CG3842, *Dhap-at*, and *MFS18*), protein glycosylation (CG11388), synaptic adhesion (*Nlg4*), a paralogue of Naa20 N-terminal acetyltransferase (CG31730), and DNA repair/transcription (*Parp*). Surprisingly, two of the strongest suppressors were proteins that participate in the SAC, Mps1 and Mad2, and whose depletion specifically suppressed *san* RNAi adult wing phenotypes (Figure 1D,E). Given that both of these proteins belong to the same biological pathway, and both were isolated as suppressors of *san* RNAi, we hypothesized that impairment of SAC could rescue mitotic defects caused by cohesin deficiency.

**Table 1.** List of genes identified in the screen.

| Type of interaction | CG | Gene <sup>1</sup> | TRiP number | Average wing phenotypic class (n) |  |
| --- | --- | --- | --- | --- | --- |
|  |  |  |  | Crossed with Nubbin-Gal4 | Crossed with Nubbin-Gal4 UAS-San RNAi |
| *Suppressors* | CG3100 | *Cad99C* | HMS01451 | 1.00 (240) | 2.35 (353) |
|  | CG3399 | *capu* | HMS00712 | 1.00 (161) | 2.40 (170) |
|  | CG4625 | *Dhap-at* | HMC03654 | 1.00 (189) | 2.49 (211) |
|  | CG3265 | *Eb1* | HMS01568 | 1.00 (175) | 2.44 (232) |
|  | CG3161 | *His3* <sup>2</sup> | GL00255 | 2.56 (223) | 2.48 (123) |
|  | CG1013 | *JMJD7* | HMS00578 | 1.00 (232) | 2.50 (263) |
|  | CG4029 | *jumu* | HMS02839 | 1.00 (186) | 2.44 (252) |
|  | CG1749 | *mad2* | GLC01381 | 1.08 (202) | 2.04 (224) |
|  | CG7643 | *Mps1* | GL00184 | 1.00 (238) | 2.22 (364) |
|  | CG3413 | *Nlg4* | HMS01710 | 1.04 (198) | 2.50 (221) |
|  | CG4041 | *Parp* | GL00229 | 1.00 (92) | 2.21 (109) |
|  | CG1173 | *Sfxn1-3* | HMS01674 | 1.00 (186) | 2.58 (361) |
|  | CG5589 | - | HMS00325 | 2.04 (257) | 2.70 (155) |
|  | CG3842 | - | HMC02378 | 1.03 (223) | 2.05 (250) |
|  | CG3173 | - | HMS02540 | 1.68 (133) | 2.24 (269) |
|  | CG1138 | - | HMS02633 | 1.00 (201) | 2.37 (336) |
|  | CG4766 | - | HMC02404 | 1.91 (204) | 2.36 (287) |
|  | CG8746 | - | HMS02863 | 1.95 (125) | 2.34 (201) |
|  | CG1259 | - | GL00498 | 1.00 (183) | 2.48 (382) |
| *Enhancers* | CG6875 | *asp* | GL00108 | 1.00 (262) | 4.49 (216) |
|  | CG4231 | *cno* | HMS00239 | 1.04 (245) | 3.81 (236) |
|  | CG3225 | *Claspin* | HMS00772 | 1.00 (168) | 4.27 (182) |
|  | CG8598 | *eco* <sup>4</sup> | GL00528 | 1.00 (191) | 4.29 (181) |
|  | GC4203 | *Mau2* <sup>4</sup> | HMS02374 | 1.00 (122) | 3.72 (182) |
|  | CG1543 | *MFS18* | HMS00961 | 1.00 (102) | 3.98 (164) |
|  | CG2163 | *Pabp2* | HMS00553 | 1.00 (153) | 3.91 (206) |
|  | CG3157 | *γTub23C* | GL01171 | 1.05 (229) | 3.54 (254) |
|  | CG9773 | - | HMS01044 | 1.11 (192) | 4.12 (124) |
|  | CG3224 | - | GL00501 | 1.00 (244) | 3.79 (254) |

<sup>1</sup> The gene abbreviations used are from Flybase version FB2017\_06.
<sup>2</sup> The RNAi used theoretically also depletes the histone H3 isoforms CG33833, CG33806, CG33839, CG33827, CG33854, CG33824, CG33818, CG33830, CG33863, CG33815, CG33866, CG33836, CG33803, CG33851, CG33809, CG33821, CG33845, CG33857, CG33848, CG33860, CG33842 and CG33812.
<sup>3</sup> This RNAi was classified as suppressor due to its strong adult wing phenotype when expressed in the wing imaginal discs with nubbin-Gal4.
<sup>4</sup> The genetic interaction with *eco* and *Mau2* has been previously described in (Ribeiro et al., 2016).

**Table 2.** Main function of the genes identified in the screen.

| CG | Gene | Human/Yeast<br>ortologue | Description | Type of<br>interaction | References<br>(PMID) |
| --- | --- | --- | --- | --- | --- |
| CG6875 | *asp* | ASPM/ - | Mitotic microtubule organization | Enhancer | 10073938 |
| CG31009 | *Cad99C* | PCDH15/ - | Cell adhesion | Suppressor | 11162110 |
| CG42312 | *cno* | AFDN/ - | Cell adhesion | Enhancer | 22024168 |
| CG3399 | *capu* | FMN2/ - | Nucleation of actin<br>microfilaments | Suppressor | 17925229 |
| CG32251 | *Claspin* | CLSPN/ - | DNA replication checkpoint | Enhancer | 22796626 |
| CG4625 | *Dhap-at* | GNPAT/ - | Peroxisomal ether lipid synthesis | Suppressor | 22758915 |
| CG3265 | *Eb1* | MAPRE1/ BIM1 | Spindle assembly, dynamics, and<br>positioning | Suppressor | 12213835 |
| CG8598 | *eco*<sup>1</sup> | ESCO1/ECO1 | Cohesion establishment | Enhancer | 14653991 |
| CG31613 | *His3* | HIST1H3B/ - | Chromatin component | Suppressor |  |
| CG10133 | *JMJD7* | JMJD7/ - | Putative histone demethylase | Suppressor |  |
| CG4029 | *jumu* | FOXN4/ - | Transcription factor | Suppressor | 10940625 |
| CG17498 | *mad2* | MAD2L1/ MAD2 | Mitotic spindle checkpoint | Suppressor | 15075237 |
| CG4203 | *Mau2*<sup>1</sup> | MAU2/ - | Cohesion loading | Enhancer | 10882066 |
| CG15438 | *MFS18* | SLC17A9/ - | Major facilitator superfamily<br>transporter | Enhancer | 22359624 |
| CG7643 | *Mps1* | TTK/ MPS1 | Spindle checkpoint | Suppressor | 15556864 |
| CG5589 | - | Ddx52/ ROK1 | ATP-dependent RNA helicase | Suppressor | 28701701 |
| CG3842 | - | -/ ENV9 | Retinoid-active short-chain<br>dehydrogenase/reductase | Suppressor | 19520149 |
| CG31730 | - | -/ - | N-terminal acetyltransferase <sup>2</sup> | Suppressor |  |
| CG11388 | - | XXYL1/ - | Protein glycosylation <sup>3</sup> | Suppressor |  |
| CG4766 | - | Mab21l2/ - | Cell fate <sup>4</sup> | Suppressor |  |
| CG8746 | - | -/ - |  | Suppressor |  |
| CG9773 | - | UNC50/ GMH1 | UNC-50 protein family | Enhancer |  |
| CG32243 | - | -/ - |  | Enhancer |  |
| CG12592 | - | SLAIN1-2/ - |  | Suppressor |  |
| CG34139 | *Nlg4* | NLGN3/ - | Synaptic adhesion protein | Suppressor | 10892652,<br>24068821 |
| CG2163 | *Pabp2* | PABPN1/ SGN1 | mRNA polyadenylation | Enhancer | 10481015 |
| CG40411 | *Parp* | PARP1/ - | DNA repair/transcription | Suppressor | 11376691,<br>24055367 |
| CG11739 | *Sfxn1-3* | SFXN1/ FSF1 | Mitochondrial ion<br>transmembrane transporter | Suppressor |  |
| CG3157 | *γTub23C* | TUBG2/ TUB4 | Microtubule organization | Enhancer | 1904010 |

<sup>1</sup> The genetic interaction with *eco* and *Mau2* has been previously (Ribeiro et al., 2016).
<sup>2</sup> Testis specific paralogue of *Drosophila* Naa20 (CG14222).
<sup>3</sup> Homologue to the human xyloside xylosyltransferase 1 that transfers the second xylose to O-glucosylated epidermal growth factor repeats of notch (Sethi et al., 2012)
<sup>4</sup> Paralogue of *Drosophila* mab-21 (CG4746).

### SAC inactivation rescues chromosome segregation defects associated with loss of cohesion

To gain further insight on whether SAC inactivation could indeed rescue cohesion defects we sought out to evaluate mitotic fidelity in various experimental conditions. Live cell imaging analysis in the developing wing disc revealed, as expected by our previous work (Ribeiro et al., 2016), that upon *san* RNAi, cells exhibited various degrees of sister chromatid cohesion defects. In control strains all cells underwent mitosis with normal metaphase morphology (Figure 2A, Figure S2 and Movie S1). Upon san RNAi only 13±10% displayed normal mitosis and most cells underwent partial or full sister chromatid separation (17±6 and 70±13%, respectively), resulting in SAC activation and extended mitosis (Figure 2A,B, Figure S2 and Movie S2). More severe defects were obtained when cohesion loss was induced by acute artificial cleavage of cohesin Rad21 subunit, using a previously established TEV protease-mediated cleavage method (Pauli et al., 2008). In these experiments, wing imaginal discs were allowed to develop normally until 3^rd^ instar larvae stage, when TEV protease was induced by heat-shock. After heat-shock Rad21 became quickly undetectable in cells expressing exclusively TEV-sensitive Rad21-EGFP (Figure S3). TEV expression resulted in full sister chromatid separation across all cells analysed (Figure S2), leading to extended mitosis and chromatid shuffling between the poles (Figure 2C,D and movie S3).

**Figure 2.**
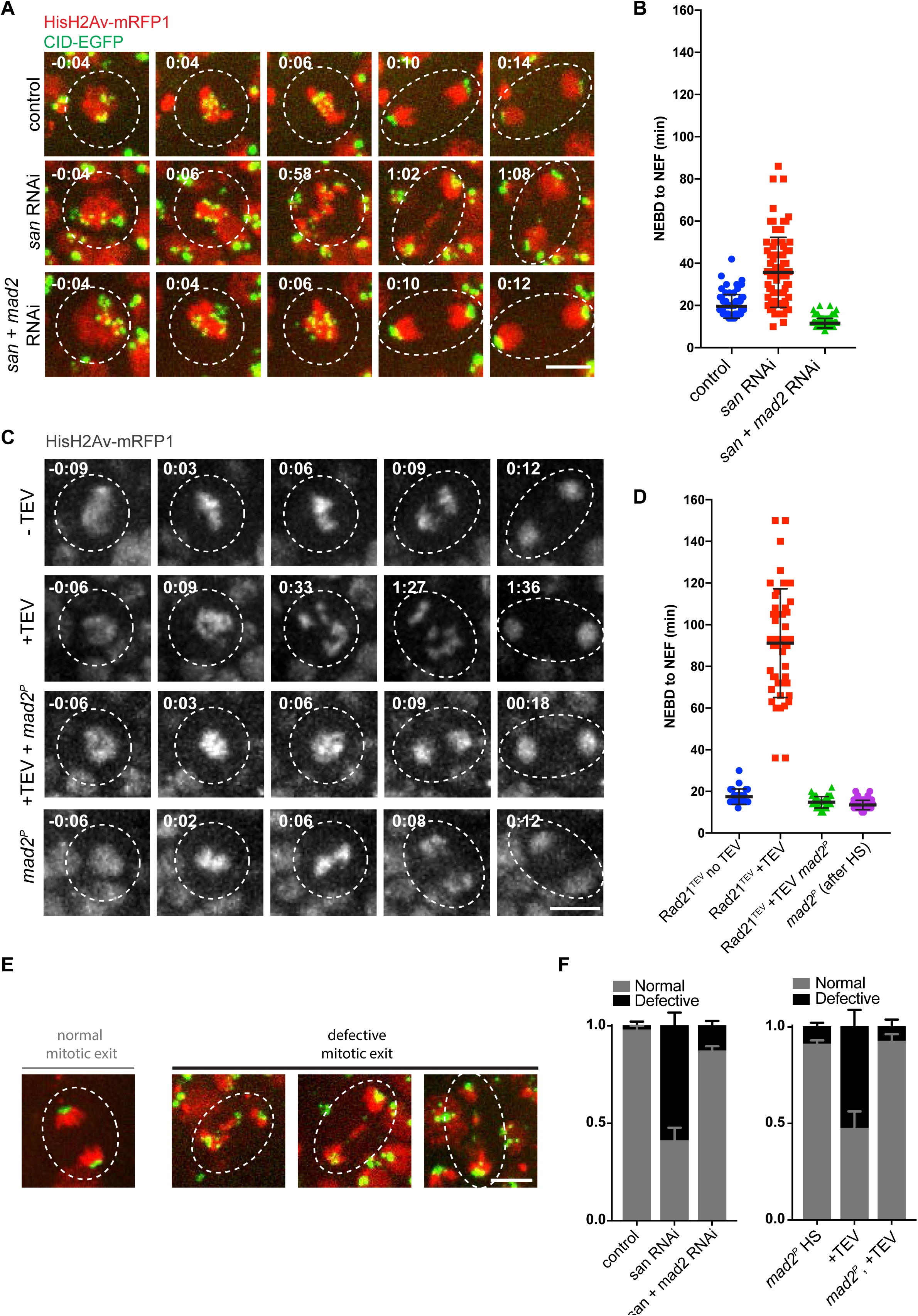
Inhibition of SAC in wing imaginal discs alleviates mitotic errors caused by premature loss of cohesin. **A.** Images from movies of the wing disc pouch in the control, *san* RNAi and *san* and *mad2* RNAi strains. Strains contained HisH2Av-RFP (red) and CID-EGFP (green). Times are relative to NEDB. Scale bar is 5 µm. **B.** Quantification of mitotic duration in control, *san* RNAi, or *san* and *mad2* RNAi strains. The duration of mitosis was measured from nuclear envelope breakdown (NEBD) to nuclear envelope formation (NEF) using H2Av-RFP channel. Images were taken every 2 minutes. Each dot represents an individual cell and lines represent mean ± SD (n= 71/5 for control, 77/5 for *san* RNAi and 124/5 for *san*+*mad2* RNAi, n=number of cells/number of independent discs). **C.** Images from movies of the wing disc from strains surviving solely on TEV-cleavable Rad21 (Rad21TEV) with and without heat-shock induced TEV protease cleavage, in strains wild type or homozygous mutant for the *mad2* gene. Strains also expressed HisH2Av-RFP (red) for visualization of mitotic duration and phenotype. Times are relative to NEDB. Scale bar is 5 µm **D.** Quantification of mitotic duration of the no heat shock control, upon TEV-protease mediated cleavage of Rad21TEV and TEV-protease mediated cleavage of Rad21^TEV^ in a *mad2* mutant background. The duration of mitosis was measured from nuclear envelope breakdown (NEBD) to nuclear envelope formation (NEF) using H2AvD-mRFP1. Images were taken every 2/3 minutes. Each dot represents an individual cell and lines represent mean ± SD (n= 27/4 for Rad21^TEV^ – TEV (no HS), 46/8 for Rad21^TEV^ + TEV, 46/4 for Rad21^TEV^+TEV in a *mad2^P^* background and 60/4 for *mad2P* after heat-shock (HS), n=number of cells/number of independent discs). **E)** Representative images of mitotic cells from *san* RNAi undergoing mitosis with normal and defective chromosome distribution. Scale bar is 5 μ*m* and applies to all images. **F)** Quantification of mitotic exit defects observed in the different experimental conditions; graph represents mean ± SEM of errors of individual discs (n≥4 independent discs corresponding to over 80 cells analyzed per experimental condition).

In order to inhibit the SAC, we focused on genetic conditions that remove Mad2, a key component of this checkpoint, as to date this protein is thought to be solely required for SAC response (in contrast to Mps1 that has been implicated in other mitotic functions (Liu and Winey, 2012). Flies carrying null alleles for the *mad2* gene were previously shown to be viable (Buffin et al., 2007) and its depletion in the larvae wing imaginal disc did not compromise wing development (Figure 1E). As expected, removal of Mad2 by RNAi or the *mad2^P^* null allele abolished the mitotic delay in both experimental conditions for cohesion loss, *san* RNAi and TEV-mediated Rad21 cleavage (Figure 2A,B,C,D and Movies S4 and S5). More importantly, shortening of mitotic timing drastically reduced the frequency of abnormal anaphase figures (Figure 2E,F). Whereas upon premature loss of cohesin mitotic exit often displays lagging chromatids or chromatin bridges, these segregation defects were significantly reduced when SAC was removed (Figure 2 E,F).

The aforementioned analysis of mitotic defects does not account for numerical errors in chromosome segregation. Therefore, a parallel evaluation of chromosome distribution symmetry was performed in early syncytial blastoderm embryos, as segregation efficiency of the synchronously dividing nuclei can be more accurately evaluated. Cohesin cleavage in *Drosophila* syncytial embryos was induced by microinjection of TEV protease during interphase, as previously described (Oliveira et al., 2010). This led to full separation of sister chromatids after NEBD and a short mitotic delay (Figure 3A,B, movies S6, S7). To test if such mitotic delay was SAC dependent, we performed similar experiments in a *mad2* mutant background. Mitotic duration under these conditions was indistinguishable from controls, implying that SAC surveillance is responsible for the delay in mitotic progression upon premature loss of sister chromatid cohesion (Figure 3A,B, movies S8). To evaluate chromosome distribution we measured the area occupied by centromeres in the vicinity of each pole as well as those lagging behind the segregation plane, ending up in the middle of the segregation plane during mitotic exit (most likely due to merotelic attachments) (Figure 3C). Segregation symmetry was evaluated as a ratio between the areas occupied by each cluster of centromeres (Cid-EGFP) during mitotic exit. As expected, this value was close to one in control strains (Figure 3D). TEV-mediated cohesin cleavage caused a high degree of asymmetry between centromeric signals placed at the poles (Figure 3D). Moreover, TEV-cleavage resulted in the presence of lagging chromosomes during anaphase (Figure 3E). Consistent with our previous results (Figure 2), loss of SAC led to a significant reduction of the segregation error frequency after TEV-cleavage, as evidenced by the significant recovery in centromere distribution symmetry and the decrease in lagging centromeres (Figure 3D,E).

**Figure 3.**
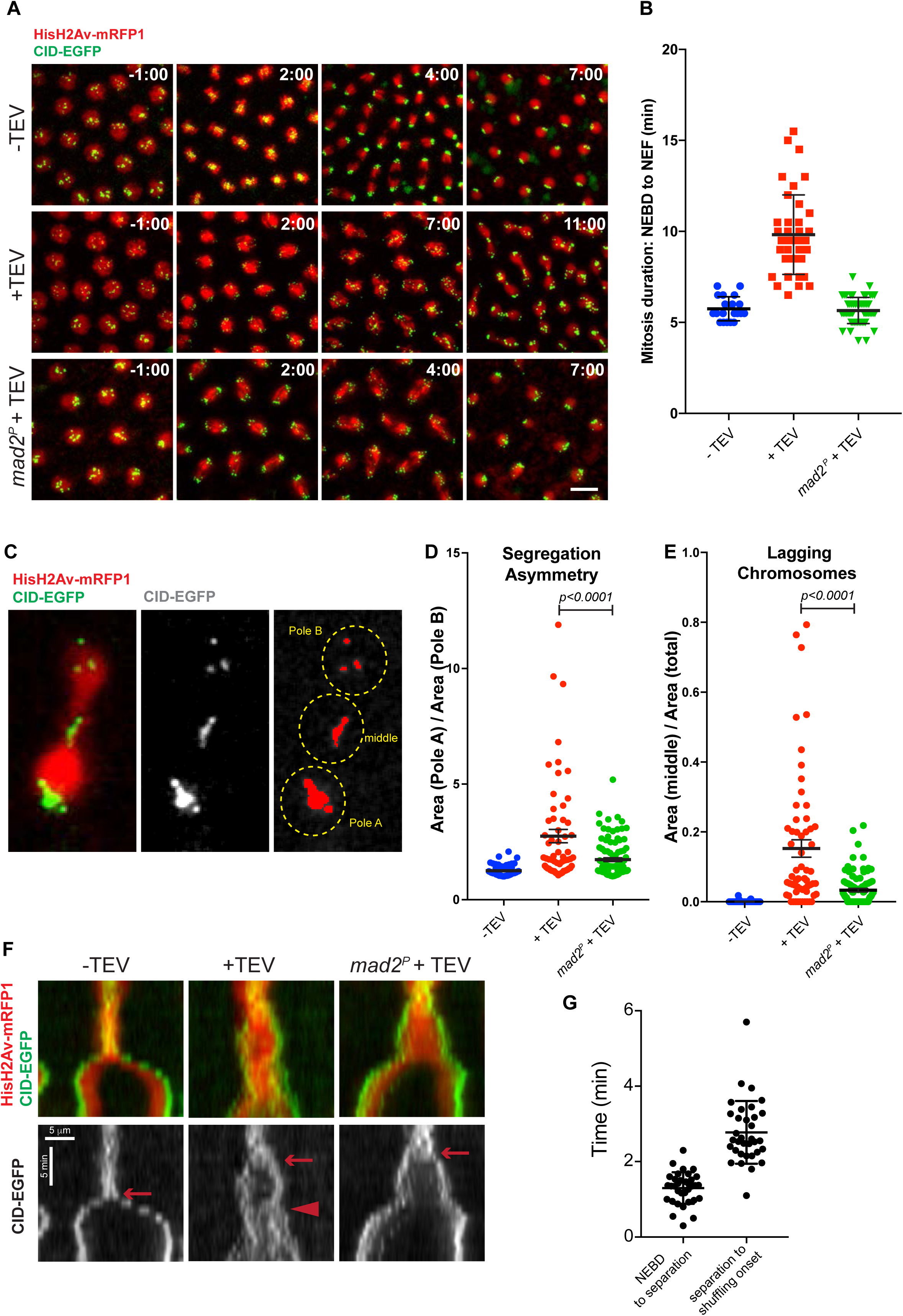
Inhibition of SAC in syncytial blastoderm embryos alleviates mitotic errors caused by premature loss of cohesin. **A.** Embryos surviving solely on Rad21^TEV^ either non-injected (up) or injected with 5 mg/ml TEV protease (middle and bottom panels). Embryos are derived from females that are wild type or homozygous mutant for *mad2* gene and express HisH2Av-RFP (red) and CID-EGFP (green). Images were taken every 30 seconds and times are relative to NEBD. Scale bar is 10 µm. **B.** Quantification of mitotic duration in un-injected embryos and embryos injected with TEV protease in strains containing solely Rad21^TEV^ and wild type or mutant for *mad2*. The duration of mitosis was measured from nuclear envelope breakdown (NEBD) to nuclear envelope formation (NEF) using HisH2Av-RFP. Images were taken every 30 seconds. Each dot represents a single mitosis and lines represent mean ± SD (n= 20/4 for Rad21^TEV^ no TEV, 40/8 for Rad21^TEV^ + TEV and 55/11 for Rad21^TEV^+TEV in a *mad2^P^* background, n=number of mitosis/number of independent embryos). **C.** Representative image from mitotic exit upon TEV-mediated cohesin cleavage, highlighting how centromere positioning was measured in D; An automatic threshold was used to determine the positioning of centromeres and the area in each pole and between segregating poles was measured. **D.** Quantification of segregation asymmetry in control, cohesin cleavage, and cohesin cleavage in *mad2* mutant background. Each value was quantified by normalizing the area of pole A (with higher area) and the area of pole B (lower area), as illustrated in C; (n= 46/5 for Rad21^TEV^ no TEV, 60/6 for Rad21^TEV^ + TEV and 60/6 for Rad21^TEV^+TEV in a *mad2^P^* background, n=number of telophases/number of independent embryos) **E.** Relative area of lagging centromeres in control, RAD21^TEV^ + TEV protease, and RAD21^TEV^ + TEV protease in a *mad2* mutant background; statistical analysis was performed using one-way ANOVA test. **F.** Kymographs of HisH2Av-RFP and CID-EGFP of cells entering mitosis in control, cohesin cleavage, and cohesin cleavage in *mad2* mutant background. Arrow points to centromere separation and arrowhead to the shuffling onset. Scale bars are 5 min and 5 µm. **G.** Quantification of time for chromosome shuffling onset upon TEV-mediated cohesin cleavage, relative to NEBD. Each dot represents a single dividing nuclei from >10 independent embryos.

Altogether, these results demonstrate that SAC inactivation rescues the chromosome segregation defects associated with premature loss of sister chromatid cohesion.

### SAC inactivation suppresses chromosome shuffling after loss of cohesion

We demonstrated that loss of SAC enhanced mitotic fidelity upon cohesin impairment. The severity of the phenotypes associated with premature cohesion loss is associated with extensive genome randomization. This is enhanced by chromosome shuffling due to engagement of isolated chromatids into error-correction reactions (Khodjakov and Pines, 2010; Mirkovic et al., 2015; Nezi and Musacchio, 2009; Oliveira et al., 2010). We postulated that SAC loss enhanced mitotic fidelity by reducing mitotic timing and thereby limiting the degree of chromosome shuffling. To evaluate this hypothesis, the timing and extent of chromosome shuffling was assessed in cells undergoing mitosis upon premature loss of sister chromatid cohesion in the absence or presence of a functional SAC. In embryos, the SAC-dependent mitotic delay observed upon cohesin cleavage, albeit short (~ 4 min) (Figure 3B), it was nevertheless long enough for a high degree of chromosome shuffling before mitotic exit (movie S7). In the absence of a functional SAC, however, and despite the evident and full premature loss of cohesion, there was a decrease in chromosome shuffling events (Figure 3A and movie S8). Similar results were obtained in wing disc cells, where chromosome-shuffling events after cohesin cleavage were infrequent upon SAC inactivation (Movies S4 and S5).

Altogether, these results suggest that despite cohesin loss, engaging into error-correction does not take place during early mitotic stages. To test this possibility, we sought out to measure the kinetics of chromosome shuffling onset upon full loss of sister chromatid cohesion. Analysis of chromosome configuration, both in embryos and wing disc cells, revealed that despite cohesin removal, chromosomes retain a pseudo-metaphase configuration for an extended period of time (Figure 2A, t=6 min; 2C, t=9min; Figure 3A t=2min; see also movies S2, S3,S7). During this pseudo-metaphase stage, sister centromeres were found fully disjoined, confirming loss of sister chromatid cohesion. However, sister chromatid individualization was only initiated later. Even upon full sister chromatid separation, there was an evident delay in the initiation of chromosome shuffling, implying that sister chromatid separation does not trigger immediate error-correction. To confirm this possibility, the timing of error-correction engagement was analysed using kymographs that plot the positioning of centromeres along the segregation plane over time. The time of centromere separation can be easily detected by the split in centromere signals and the onset of chromosome shuffling by the time centromeres start crossing the middle of the segregation plane (Figure 3F arrow and arrow heads, respectively). This analysis revealed that upon cohesin cleavage, chromosome shuffling was only initiated 4.07± 0.96 min after NEBD (1.3±0.4min for NEBD to centromere separation and 2.8±0.8min from separation to initiation of shuffling) (Figure 3G). This analysis reveals a significant delay in the initiation of major error-correction events. A similar, yet extended behaviour, was also observed in larvae wing disc cells. Upon NEBD, chromosomes retained a prolonged pseudo-metaphase configuration despite sister chromatid separation (as judged by centromere distances) and chromosome shuffling was only observed much later (11.4± 2.9min after NEBD, Figure S4). The observed delay in extensive shuffling engagement is similar to the mitotic timing in the absence of a functional SAC (Figures 2B,D and 3B). Thus, SAC counteracts genome shuffling in the absence of cohesin by shortening mitosis duration and thereby preventing extensive error-correction.

### SAC inactivation restores cell survival after loss of cohesion

Our results indicate that the mitotic defects upon loss of cohesion are less detrimental in the absence of SAC. If so, the degree of aneuploidy should follow a similar trend. Larvae wing discs are well known to eliminate cells with an erroneous DNA content by apoptosis (Dekanty et al., 2012; Poulton et al., 2014). Therefore, SAC inactivation should reduce the levels of apoptosis after loss of cohesion. To evaluate the extent of apoptosis upon full cohesion loss, larvae carrying a TEV-sensitive Rad21 were heat-shocked to induce TEV protease expression and wing discs were dissected 24h after heat-shock. Virtually no apoptosis was detected (staining for cleaved caspase 3 (CC3)) within the control wing discs (Rad21^TEV^ in the absence of TEV protease) (Figure 4A,B). In contrast, TEV-mediated cohesin cleavage induced high levels of apoptosis within 24hr, extending to over 15±2% (mean ± SD, n=7) of the entire wing disc area (Figure 4A,B). Remarkably, the levels of apoptosis were significantly reduced if cohesin loss was induced in the absence of a functional SAC (3±3%, mean ± SD, n=10) (Figure 4A,B). Similar results were obtained upon depletion of San (*san* RNAi), where apoptosis covered approximately 7±4% (mean ± SD, n=5) of the wing disc pouch area, compared to only approximately 0.9±0.6% (mean ± SD, n=6) of the pouch area after co-depletion of San and Mad2 (Figure 4C,D). These results show that inactivation of SAC in a proliferating tissue significantly increases cell survival upon loss of cohesion.

**Figure 4.**
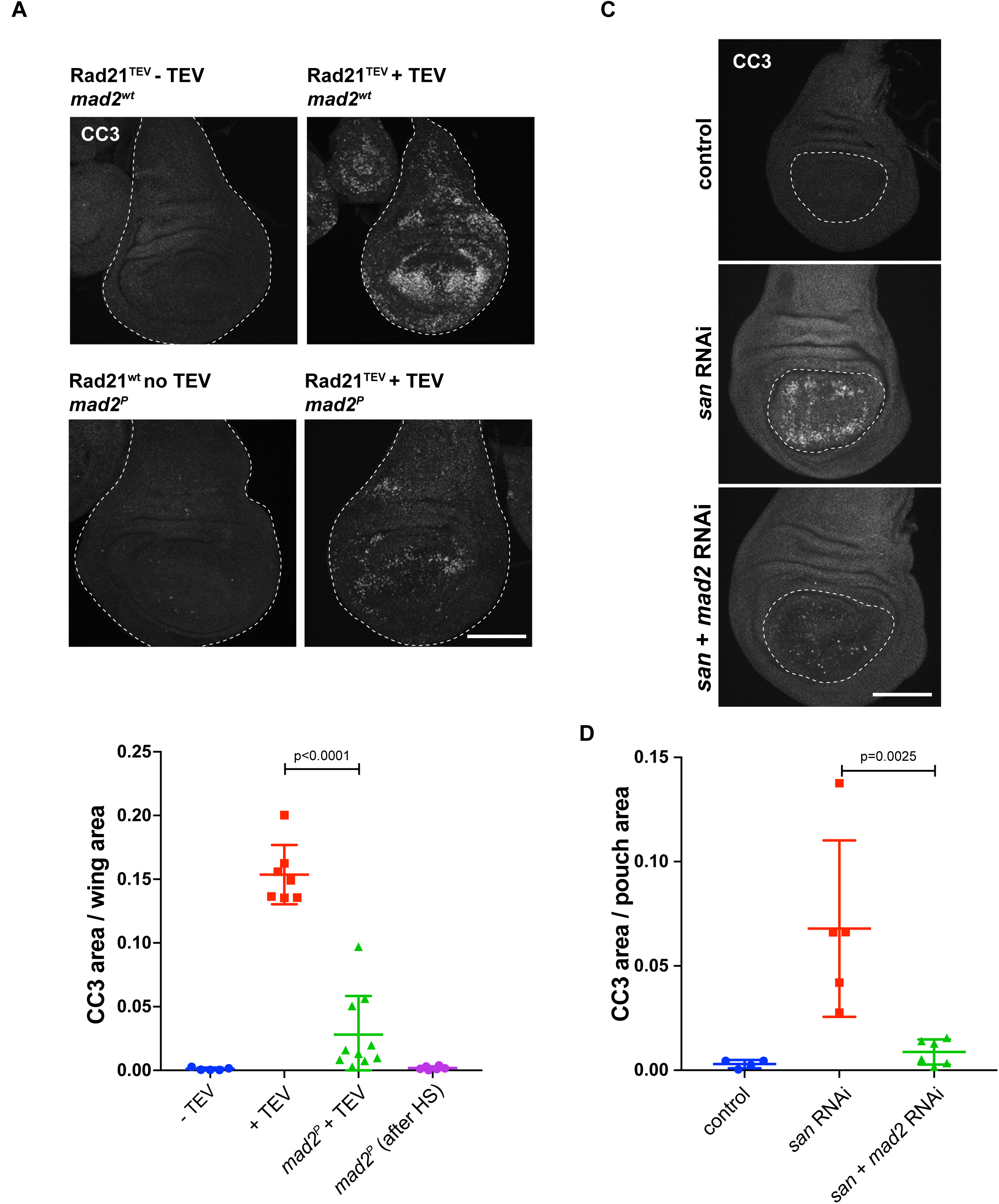
Inhibition of SAC suppresses imaginal wing disc apoptosis caused by premature loss of cohesin. **A.** Images of Cleaved Caspase 3 (CC3) immunofluorescence in controls (Rad21^TEV^ without TEV and *mad2^P^* after heat-shock (HS)), RAD21^TEV^ + TEV protease and RAD21^TEV^ + TEV protease in *mad2* mutant background after HS. Scale bar is 100 µm. **B.** Quantification of CC3 positive area of the entire wing disc, in the indicated experimental conditions; n≥ 5 independent discs per experimental condition; statistical analysis was performed using one-way ANOVA test. **C.** Representative images of CC3 immunofluorescence in control, *san* RNAi and *san* and *mad2* double RNAi. **D.** Quantification of CC3 positive area of the wing disc pouch, in control, *san*, and *san* and *mad2* RNAi; n≥ 4 independent discs per experimental condition; multiple comparison analysis was perform using a one-way ANOVA test.

## Discussion

In agreement with the “safeguard” function for the Spindle Assembly Checkpoint, mitotic errors are often exacerbated by impairment of the SAC. These include defects associated with multiple centrosomes, defective microtubule assembly or kinetochore structure (Daniel et al., 2006; Gogendeau et al., 2015; Lee and Spencer, 2004; Poulton et al., 2017; Tarailo et al., 2007). Here we demonstrate that the opposite happens with regard to cohesion defects. Absence of the SAC alleviated mitotic errors and improved mitotic fidelity after cohesion loss. Cells with a functional SAC undergo extensive chromosome shuffling and consequent randomization of the genome, whereas virtually no shuffling could be observed in absence of the SAC. The detrimental nature of SAC in the presence of cohesion defects is likely related to the irreversibility of cohesion loss. Most mitotic defects can be corrected over time (e.g. SAC-mediated mitotic delay enables clustering of multiple centrosomes (Basto et al., 2008)). In sharp contrast, premature cohesin loss is an irreversible error and prolonging mitosis duration further enhances genome randomization.

The improved mitotic fidelity after cohesion loss in the absence of SAC is likely a consequence of slow kinetics of error-correction engagement coupled with a bias for chromosome orientation towards a correct alignment. Several mechanisms are known to bias chromosome segregation towards the right orientation, including centromere geometry (Tanaka et al., 2000), bias in microtubule growth towards the kinetochores (Carazo-Salas et al., 1999; Wollman et al., 2005) and/or kinetochore-mediated microtubule nucleation (Kitamura et al., 2010; Maiato et al., 2004). Of these, chromosome geometry is believed to facilitate bipolar attachment by facing one kinetochore to the opposite pole upon attachment to a pole. If so, what ensures geometric arrangement during the initial mitotic stages, even in the absence of cohesin? A possible mechanism enabling a transient organization of sister chromatids towards opposing poles is incomplete resolution of sister chromatid intertwines. Yet, residual catenation present in metaphase chromosomes is unable to confer functional cohesion as removal of cohesin is sufficient to induce immediate sister chromatid separation (Oliveira et al., 2010; Uhlmann et al., 2000). Additional mechanisms may thus impair prompt resolution of sister chromatids specifically during early mitosis, in contrast what is observed in metaphase chromosomes. Spindle forces were described to enhance decatenation (Baxter et al., 2011; Charbin et al., 2014; Mariezcurrena and Uhlmann, 2017) and thus resolution of several DNA-intertwines may only be achieved upon chromosome capture. Recent findings propose that efficient decatenation requires constant “guiding action” from condensin I (Piskadlo et al., 2017). Maximal levels of this complex are only observed on mitotic chromosomes once in late-metaphase/anaphase (Gerlich et al., 2006; Oliveira et al., 2007), which could also limit full decatenation to the later stages of mitosis. Residual catenation is unable to sustain cohesion and chromosome alignment in a SAC competent cell. Yet, it may be sufficient to allow a transient pseudo-metaphase alignment that biases initial chromosome attachment to the right orientation. This is certainly an error-prone process but nevertheless more accurate when compared to total genome randomization due to extensive chromosome shuffling.

Why separated single sisters are inefficient at triggering error-correction mechanisms during early mitosis remains to be addressed. This could be related to a partial tension state facilitated by pseudo-metaphase chromosomal configuration, precluding error-correction activation. Additionally, an intrinsic delayed action of error correction machinery may further account for observed late shuffling onset. Indeed, slow kinetics or a lag time of Aurora B-mediated chromosome detachment has been hypothesized in several theoretical studies (Kalantzaki et al., 2015; Tubman et al., 2017; Zhang et al., 2013) but so far little experimental observations support this claim. Such intrinsic delay would solve the “problem of initiation of biorientation” whereby initial interactions (necessarily under low tension) are able to survive such a tension-sensitive mechanism for chromosome detachment (Khodjakov and Pines, 2010; Tubman et al., 2017).

Interestingly, the interplay between mitotic timing and sister chromatid cohesion has been previously reported in mammalian cells whereby extension of mitosis predisposes to sister chromatid cohesion defects. Cells arrested in mitosis for long periods were shown to display sister chromatid separation (referred as “cohesion fatigue”) (Daum et al., 2011). Moreover, defective sister chromatid cohesion was described to be synthetically lethal with impaired APC/C function in Warsaw breakage syndrome (WABS) patient-derived cells as well as several cancer cell lines with cohesion defects (de Lange et al., 2015). Our observations now demonstrate how reduction of mitotic timing is sufficient to rescue segregation defects associated with premature cohesin loss. Importantly, these experiments highlight the detrimental effect of the SAC upon cohesion defects. When sister chromatid cohesion is compromised, and thus mitotic fidelity irreversibly affected, the SAC exacerbates mitotic errors in contrast to its canonical protective function.

## Methods

### *Drosophila* strains and rearing conditions

*Drosophila melanogaster* flies were raised at 25ºC or 18ºC for hs-TEV containing crosses in polypropylene vials (51 mm diameter) containing enriched medium (cornmeal, molasses, yeast, soya flour and beetroot syrup). All RNAi lines used in the screen (Table S1) are from the Transgenic RNAi project (TRiP) and are available in the Bloomington Drosophila Stock Center. Other *Drosophila* stocks used in this study are indicated in Table S5. To induce full cohesin cleavage in a temporally controlled manner, by TEV protease cleavage, *Drosophila* strains were used with TEV-cleavable Rad21 (Rad21^TEV^) in a Rad21-null background (*rad21^ex15^, Rad21^271-3TEV^-myc* or *rad21^ex15^, Rad21^550-3TEV^-EGFP*) (Oliveira et al., 2014; Pauli et al., 2008), in strains mutant or wild type for the Mad2 gene (Althoff et al., 2012; Buffin et al., 2007). TEV expression was induced by heat-shocking 3^rd^ instar larvae at 37ºC for 45 minutes. Larvae were then left to recover at room temperature. For live cell imaging, fly strains also expressed *His2AvD-mRFP1* and *Cid-EGFP* (Schuh et al., 2007) fluorescent markers.

### *Drosophila* screen Details

In the screen we analysed 2955 RNAi lines that theoretically deplete 2920 proteins, corresponding to approximately 21% of all protein coding genes annotated in Flybase (Flybase versionFB2017_06). To select the lines to test we followed a list of available RNAi downloaded from the TRiP website. In this list, the lines are ordered alphabetically, according to gene name or CG number. However, our results do not strictly follow this list, since we mainly used lines constructed with Vallium20 or Valium22 vectors and some lines did not survive shipping. In the screen, females carrying the nubbin-Gal4, UAS-*san* RNAi were crossed with males of different RNAi lines from TRiP (see diagram in Figure S1). The progeny of these flies were classified into different classes according to the adult wing phenotypes: class 1 - wild type wings; class 2 – flies with wings that present only mild morphological defects; class 3- flies whose wing morphological defects are intermediate (similar *san* RNAi); class 4 – flies whose wings show strong morphological defects; class 5 – flies without wings or vestigial wings (Figure 1C) (Wu and Cohen, 2002). The average adult wing class for each condition was always calculated using more than 50 adult flies (n≥50). If the average class for a given genetic interaction was equal or below 2.6 than the RNAi line tested was classified as suppressor, if the average class was equal or above 3.5 than the RNAi line was classified as enhancer (Figure S1). To exclude RNAi lines whose expression by itself led to wing morphological defects, in otherwise wild type imaginal discs, we crossed all lines carrying RNAis identified in the first cross with nubbin-Gal4 and discarded all RNAi lines that were enhancers and produced significant phenotypes by itself (Figure S1).

### Live imaging

For imaging of wing discs, larval imaginal wing discs were dissected in Schneider medium with 10% FBS. Dissected discs were placed and oriented in a 200 *μ* l drop of medium at the bottom of a glass-bottom petridish (MakTek). Time lapse imaging analysis was performed on a spinning disc microscope using either a Revolution XD microscope (Andor, UK), equipped with immersion a 60x (water) and 100x (oil) objectives (Nikon, Japan) and a iXon +512 EMCCD camera (Andor, UK), or a Revolution XD microscope (Andor, UK) equipped with immersion a 60x glycerol-immersion 1.30 NA objective (Leica Microsystems, Germany) and a 100x oil-immersion 1.4 NA objective (Leica Microsystems, Germany) and a iXon Ultra 888 1024*1024 EMCCD (Andor, UK). Stacks of 20-30 frames 0,5 *μ* m apart were taken every 1 to 3 minutes. For syncytial embryo imaging, embryos were aligned on coverslips and covered with Series 700 halocarbon oil (Sigma-Aldrich). Time-lapse microscopy was performed with an inverted wide-field DeltaVision microscope (Applied Precision Inc., Issaquah, WA) in a temperature-controlled room (18–20°C). One stack of 15 frames (0.8 mm apart) was acquired every 30 sec with a 100x 1.4 oil immersion objective (Olympus, Japan) and captured by an EMCCD camera (Roper Cascade 1024, Roper Technologies). Movies were assembled using FIJI software (Schindelin et al., 2012) and selected stills were processed with Photoshop CS6 (Adobe).

### Microinjections

Microinjections were performed as previously described (Oliveira et al., 2010; Piskadlo et al., 2017). 1-1.5 hr old embryos were collected and processed according to standard protocols, and were injected at the posterior pole. Injections were performed using a Burleigh Thorlabs Micromanipulator, a Femtojet microinjection system (Eppendorf, Germany), and pre-pulled Femtotip I needles (Eppendorf). TEV protease was injected at 5 mg/ml TEV protease in 20 mM Tris-HCl at pH 8.0, 1 mM EDTA, 50 mM NaCl and 2 mM DTT.

### Immunofluorescence

Third instar wing imaginal disc fixation and staining was performed using standard procedures (Lee and Treisman, 2001). Briefly, third instar larvae wing disc tissue (still attached to the larva body) was fixed on ice for 30 min. The fixative consisted of 4% formaldehyde (Polysciences) in 1X PEM buffer solution. Following were washed by gentle agitation three times for 20 min in PBS-T (1x PBS + 0.1% Triton X-100). Primary antibodies incubation was performed overnight at 4 °C in PBS-T supplemented with 1% BSA and 1% donkey serum. The following day, the tissues were washed again and incubated for 2h at room temperature with the appropriate secondary antibodies diluted in PBS-T solution. Finally, after the wash of the secondary antibodies, wing discs were mounted in Vectashield (Vector Laboratories). Fluorescence images were acquired with a ×40 HCX PL APO CS oil immersion objective (numerical aperture: 1.25–0.75) on a Leica SP5 confocal microscope. Rabbit anti-cleaved caspase 3 at 1:300 (Cell Signaling, 9661S) and anti-Rabbit Alexa Fluor 488 at 1:1000 (Molecular Probes).

### Quantifications and statistics

Imaging analysis was performed using FIJI software (Schindelin et al., 2012). Statistical analysis and graphic representations were performed using Prism 7 software. Multiple comparisons were performed using one-way ANOVA, using the Bonferroni’s multiple comparison test. Graphs depict mean ± standard deviations (SD) or mean ± standard error of the mean (SEM), as indicated. Sample size details are included in the respective figure legends.

## Acknowledgements

We thank S. Heidmann, C. Lehner, R. Karess and the Bloomington Stock Center for fly strains and antibodies, all the members of the Oliveira and Martinho laboratories for discussions, Ricardo Matos for assistance with graphic design and Bárbara Kellen for technical assistance in the pilot screen. We acknowledge the TRiP at Harvard Medical School (NIH/NIGMS R01-GM084947) for providing several transgenic RNAi fly stocks used in this study. The following authors were supported by Portuguese national funding (Fundação para a Ciência e Tecnologia, FCT), fellowships: Rui D. Silva (SFRH/BPD/87482/2012), Mihailo Mirkovic (SFRH /BD/52438/2013) and Om S. Rathore (PD/BD/52428/2013, within the scope of the ProRegeM PhD program Ref. PD/00117/2012, CRM:0027030). Rui Gonçalo Martinho is supported by funding from the Association for International Cancer Research [AICR 10–0553] and the following FCT grants: PTDC/BEX-BID/0395/2014 and UID/BIM/04773/2013 CBMR 1334. Raquel A Oliveira is supported by the following grants: FCT Investigator grant (IF/00851/2012/CP0185/CT0004), EMBO Installation Grant (IG2778) and European Research Council Starting Grant (ERC-2014-STG-638917).

## Authors Contributions

R.D.S.: Conceptualization, Investigation, Writing (review & editing). M.M.: Conceptualization, Investigation, Writing (review & editing). L.G.G.: Investigation. O.S.R.: Investigation. R.G.M: Conceptualization, Writing (original draft + review & editing), Funding acquisition. R.A.O.: Conceptualization, Investigation, Writing (original draft + review & editing), Funding acquisition.

## References

Althoff, F., R.E. Karess, and C.F. Lehner. 2012. Spindle checkpoint-independent inhibition of mitotic chromosome segregation by Drosophila Mps1. Mol Biol Cell. 23:2275–2291.

Basto, R., K. Brunk, T. Vinadogrova, N. Peel, A. Franz, A. Khodjakov, and J.W. Raff. 2008. Centrosome amplification can initiate tumorigenesis in flies. Cell. 133:1032–1042.

Baxter, J., N. Sen, V.L. Martinez, M.E. De Carandini, J.B. Schvartzman, J.F. Diffley, and L. Aragon. 2011. Positive supercoiling of mitotic DNA drives decatenation by topoisomerase II in eukaryotes. Science. 331:1328–1332.

Brand, A.H., and N. Perrimon. 1993. Targeted gene expression as a means of altering cell fates and generating dominant phenotypes. Development. 118:401–415.

Buffin, E., D. Emre, and R.E. Karess. 2007. Flies without a spindle checkpoint. Nature cell biology. 9:565–572.

Carazo-Salas, R.E., G. Guarguaglini, O.J. Gruss, A. Segref, E. Karsenti, and I.W. Mattaj. 1999. Generation of GTP-bound Ran by RCC1 is required for chromatin-induced mitotic spindle formation. Nature. 400:178–181.

Charbin, A., C. Bouchoux, and F. Uhlmann. 2014. Condensin aids sister chromatid decatenation by topoisomerase II. Nucleic Acids Res. 42:340–348.

Daniel, J.A., B.E. Keyes, Y.P. Ng, C.O. Freeman, and D.J. Burke. 2006. Diverse functions of spindle assembly checkpoint genes in Saccharomyces cerevisiae. Genetics. 172:53–65.

Daum, J.R., T.A. Potapova, S. Sivakumar, J.J. Daniel, J.N. Flynn, S. Rankin, and G.J. Gorbsky. 2011. Cohesion fatigue induces chromatid separation in cells delayed at metaphase. Current biology : CB. 21:1018–1024.

de Lange, J., A. Faramarz, A.B. Oostra, R.X. de Menezes, I.H. van der Meulen, M.A. Rooimans, D.A. Rockx, R.H. Brakenhoff, V.W. van Beusechem, R.W. King, J.P. de Winter, and R.M. Wolthuis. 2015. Defective sister chromatid cohesion is synthetically lethal with impaired APC/C function. Nature communications. 6:8399.

Dekanty, A., L. Barrio, M. Muzzopappa, H. Auer, and M. Milan. 2012. Aneuploidy-induced delaminating cells drive tumorigenesis in Drosophila epithelia. Proceedings of the National Academy of Sciences of the United States of America. 109:20549–20554.

Foley, E.A., and T.M. Kapoor. 2013. Microtubule attachment and spindle assembly checkpoint signalling at the kinetochore. Nature reviews. Molecular cell biology. 14:25–37.

Gerlich, D., T. Hirota, B. Koch, J.M. Peters, and J. Ellenberg. 2006. Condensin I Stabilizes Chromosomes Mechanically through a Dynamic Interaction in Live Cells. Current biology : CB. 16:333–344.

Gogendeau, D., K. Siudeja, D. Gambarotto, C. Pennetier, A.J. Bardin, and R. Basto. 2015. Aneuploidy causes premature differentiation of neural and intestinal stem cells. Nature communications. 6:8894.

Guacci, V., D. Koshland, and A. Strunnikov. 1997. A direct link between sister chromatid cohesion and chromosome condensation revealed through the analysis of MCD1 in S. cerevisiae. Cell. 91:47–57.

Haering, C.H., A.M. Farcas, P. Arumugam, J. Metson, and K. Nasmyth. 2008. The cohesin ring concatenates sister DNA molecules. Nature. 454:297–301.

Hou, F., C.W. Chu, X. Kong, K. Yokomori, and H. Zou. 2007. The acetyltransferase activity of San stabilizes the mitotic cohesin at the centromeres in a shugoshin-independent manner. The Journal of cell biology. 177:587–597.

Kalantzaki, M., E. Kitamura, T. Zhang, A. Mino, B. Novak, and T.U. Tanaka. 2015. Kinetochore-microtubule error correction is driven by differentially regulated interaction modes. Nature cell biology. 17:530.

Khodjakov, A., and J. Pines. 2010. Centromere tension: a divisive issue. Nature cell biology. 12:919–923.

Kitamura, E., K. Tanaka, S. Komoto, Y. Kitamura, C. Antony, and T.U. Tanaka. 2010. Kinetochores generate microtubules with distal plus ends: their roles and limited lifetime in mitosis. Developmental cell. 18:248–259.

Lee, M.S., and F.A. Spencer. 2004. Bipolar orientation of chromosomes in Saccharomyces cerevisiae is monitored by Mad1 and Mad2, but not by Mad3. Proceedings of the National Academy of Sciences of the United States of America. 101:10655–10660.

Liu, X., and M. Winey. 2012. The MPS1 family of protein kinases. Annu Rev Biochem. 81:561–585.

Maiato, H., C.L. Rieder, and A. Khodjakov. 2004. Kinetochore-driven formation of kinetochore fibers contributes to spindle assembly during animal mitosis. The Journal of cell biology. 167:831–840.

Mariezcurrena, A., and F. Uhlmann. 2017. Observation of DNA intertwining along authentic budding yeast chromosomes. *Genes Dev*.

Michaelis, C., R. Ciosk, and K. Nasmyth 1997. Cohesins: chromosomal proteins that prevent premature separation of sister chromatids. Cell. 91:35–45.

Mirkovic, M., L.H. Hutter, B. Novak, and R.A. Oliveira. 2015. Premature Sister Chromatid Separation Is Poorly Detected by the Spindle Assembly Checkpoint as a Result of System-Level Feedback. Cell reports. 13:470–478.

Mirkovic, M., and R.A. Oliveira. 2017. Centromeric Cohesin: Molecular Glue and Much More. Prog Mol Subcell Biol. 56:485–513.

Musacchio, A., and E.D. Salmon. 2007. The spindle-assembly checkpoint in space and time. Nature reviews. Molecular cell biology. 8:379–393.

Nezi, L., and A. Musacchio. 2009. Sister chromatid tension and the spindle assembly checkpoint. Curr Opin Cell Biol. 21:785–795.

Ng, M., F.J. Diaz-Benjumea, J.P. Vincent, J. Wu, and S.M. Cohen. 1996. Specification of the wing by localized expression of wingless protein. Nature. 381:316–318.

Oliveira, R.A., R.S. Hamilton, A. Pauli, I. Davis, and K. Nasmyth. 2010. Cohesin cleavage and Cdk inhibition trigger formation of daughter nuclei. Nature cell biology. 12:185–192.

Oliveira, R.A., S. Heidmann, and C.E. Sunkel. 2007. Condensin I binds chromatin early in prophase and displays a highly dynamic association with Drosophila mitotic chromosomes. Chromosoma. 116:259–274.

Oliveira, R.A., S. Kotadia, A. Tavares, M. Mirkovic, K. Bowlin, C.S. Eichinger, K. Nasmyth, and W. Sullivan. 2014. Centromere-independent accumulation of cohesin at ectopic heterochromatin sites induces chromosome stretching during anaphase. PLoS biology. 12:e1001962.

Pauli, A., F. Althoff, R.A. Oliveira, S. Heidmann, O. Schuldiner, C.F. Lehner, B.J. Dickson, and K. Nasmyth. 2008. Cell-type-specific TEV protease cleavage reveals cohesin functions in Drosophila neurons. Developmental cell. 14:239–251.

Piskadlo, E., A. Tavares, and R.A. Oliveira. 2017. Metaphase chromosome structure is dynamically maintained by condensin I-directed DNA (de)catenation. Elife. 6.

Poulton, J.S., J.C. Cuningham, and M. Peifer. 2014. Acentrosomal Drosophila epithelial cells exhibit abnormal cell division, leading to cell death and compensatory proliferation. Developmental cell. 30:731–745.

Poulton, J.S., J.C. Cuningham, and M. Peifer. 2017. Centrosome and spindle assembly checkpoint loss leads to neural apoptosis and reduced brain size. The Journal of cell biology. 216:1255–1265.

Ribeiro, A.L., R.D. Silva, H. Foyn, M.N. Tiago, O.S. Rathore, T. Arnesen, and R.G. Martinho. 2016. Naa50/San-dependent N-terminal acetylation of Scc1 is potentially important for sister chromatid cohesion. Sci Rep. 6:39118.

Rong, Z., Z. Ouyang, R.S. Magin, R. Marmorstein, and H. Yu. 2016. Opposing Functions of the N-terminal Acetyltransferases Naa50 and NatA in Sister-chromatid Cohesion. J Biol Chem. 291:19079–19091.

Schindelin, J., I. Arganda-Carreras, E. Frise, V. Kaynig, M. Longair, T. Pietzsch, S. Preibisch, C. Rueden, S. Saalfeld, B. Schmid, J.Y. Tinevez, D.J. White, V. Hartenstein, K. Eliceiri, P. Tomancak, and A. Cardona. 2012. Fiji: an open-source platform for biological-image analysis. Nat Methods. 9:676–682.

Schuh, M., C.F. Lehner, and S. Heidmann. 2007. Incorporation of Drosophila CID/CENP-A and CENP-C into Centromeres during Early Embryonic Anaphase. Current biology : CB. 17:237–243.

Sethi, M.K., F.F. Buettner, A. Ashikov, V.B. Krylov, H. Takeuchi, N.E. Nifantiev, R.S. Haltiwanger, R. Gerardy-Schahn, and H. Bakker. 2012. Molecular cloning of a xylosyltransferase that transfers the second xylose to O-glucosylated epidermal growth factor repeats of notch. J Biol Chem. 287:2739–2748.

Tanaka, T., J. Fuchs, J. Loidl, and K. Nasmyth. 2000. Cohesin ensures bipolar attachment of microtubules to sister centromeres and resists their precocious separation. Nature cell biology. 2:492–499.

Tarailo, M., S. Tarailo, and A.M. Rose. 2007. Synthetic lethal interactions identify phenotypic “interologs” of the spindle assembly checkpoint components. Genetics. 177:2525–2530.

Tubman, E.S., S. Biggins, and D.J. Odde. 2017. Stochastic Modeling Yields a Mechanistic Framework for Spindle Attachment Error Correction in Budding Yeast Mitosis. Cell Syst. 4:645–650 e645.

Uhlmann, F., F. Lottspeich, and K. Nasmyth. 1999. Sister-chromatid separation at anaphase onset is promoted by cleavage of the cohesin subunit Scc1. Nature. 400:37–42.

Uhlmann, F., D. Wernic, M.A. Poupart, E.V. Koonin, and K. Nasmyth. 2000. Cleavage of cohesin by the CD clan protease separin triggers anaphase in yeast. Cell. 103:375–386.

Williams, B.C., C.M. Garrett-Engele, Z. Li, E.V. Williams, E.D. Rosenman, and M.L. Goldberg. 2003. Two putative acetyltransferases, san and deco, are required for establishing sister chromatid cohesion in Drosophila. Current biology : CB. 13:2025–2036.

Wollman, R., E.N. Cytrynbaum, J.T. Jones, T. Meyer, J.M. Scholey, and A. Mogilner. 2005. Efficient chromosome capture requires a bias in the ‘search-and-capture’ process during mitotic-spindle assembly. Current biology : CB. 15:828–832.

Wu, J., and S.M. Cohen. 2002. Repression of Teashirt marks the initiation of wing development. Development. 129:2411–2418.

Zhang, T., R.A. Oliveira, B. Schmierer, and B. Novak. 2013. Dynamical scenarios for chromosome bi-orientation. Biophysical journal. 104:2595–2606.

